# A high-resolution subcellular map of proteins in cells with motile cilia

**DOI:** 10.1101/2025.03.18.643967

**Authors:** Filippa Bertilsson, Feria Hikmet, Jan N. Hansen, Mathias Uhlén, Loren Méar, Cecilia Lindskog

## Abstract

Motile cilia are complex structures regulated by thousands of genes, essential for various physiological functions like respiration and reproduction. Their dysfunction can result in severe conditions like primary ciliary dyskinesia (PCD), highlighting the need for a deeper molecular understanding of their specific ciliary compartments. Interestingly, ciliated cells harbor multiple proteins with limited evidence on biological function, as defined by Functional Evidence (FE) scores, a grading system developed by the Human Proteome Project (HPP). Building upon the stringent antibody validation pipeline of the Human Protein Atlas (HPA) project, we developed a high-throughput workflow that combines a novel multiplex immunohistochemistry protocol with image analysis to investigate protein expression and subcellular localization in motile ciliated cells across five human tissues: nasopharynx, bronchus, fallopian tube, endometrium, and cervix. We spatially mapped >180 proteins, out of which 73% have FE scores 2-5, suggesting that further evidence is needed to establish these proteins’ biological function. Notably, expression patterns varied between tissues, suggesting that motile cilia proteins are not universally expressed across the different epithelia. Our pipeline constitutes a promising resource for comprehensive mapping of the motile cilia proteome, and a first step towards identifying cilia proteins for functional studies to understand the molecular mechanisms underlying ciliopathies.

## Introduction

Cells are not organized randomly, but instead form complex organ systems where molecules, cellular compartments and cell types are spatially distributed based on cellular function. Proteins often execute their functions in specific subcellular locations, which is crucial for maintaining various cellular processes such as cell growth, motility, and signaling1. To investigate the role of proteins and their biochemical functions within the cell, knowledge about the spatial localization from both a cellular and subcellular perspective is a crucial first step for further functional studies2.

The Human Proteome Project (HPP) is an initiative of the Human Proteome Organization (HUPO) that has contributed to credibly identifying at least one major isoform of all human proteins, mainly using mass spectrometry-based methods3. As a next step, the HPP consortium currently leads a major effort called the “Grand Challenge”, with the ultimate goal to gain a solid understanding of at least one biological function for each protein at the molecular level. A Functional Evidence (FE) scoring system has been developed, with FE1-5 elucidating the amount of evidence for protein function. The scoring system is linked to annotations contained in UniProt4, including Gene Ontology terms, publications and computational predictions where FE1 represent proteins with at least one known molecular function based on experimental evidence. FE5 represents the lowest scoring, indicating that basically nothing is known about the role of these proteins in human cells. Interestingly, only 5,229 proteins encoded by the human genome currently have FE1, highlighting the importance of further efforts that can contribute to understanding protein function.

With the goal of mapping all human proteins, the Human Protein Atlas (HPA) project has built one of the world’s largest biological knowledge resources, available as an open-access database (www.proteinatlas.org). The HPA integrates both transcriptomics data and spatial antibody-based proteomics5, and within this effort, >15,000 proteins have been mapped to all major human normal and cancer tissues using classical immunohistochemistry (IHC). The resource of more than 10 million manually annotated high-resolution histological images provides important information on spatial expression patterns of proteins from a body-wide and cell type-specific perspective, but the possibilities to distinguish subcellular patterns are limited.

Standard IHC is primarily used to study one protein per tissue section, but an increasing number of methods utilize multiplexed IHC (mIHC), enabling novel insights into co-expression and tissue or cell type heterogeneity. Spatial proteomics technologies are of utmost importance to characterize the intrinsic subcellular functions of human proteins6, as has been demonstrated in multiple large-scale studies2,7. To increase the resolution of the Tissue resource, the HPA version 24 and onwards includes data based on a novel mIHC protocol employing in-house generated 6-plex antibody panels. Using an iterative fluorescence-based staining-stripping method, fixed antibody panels of five markers outlining specific cell types, cell states or subcellular structures were stained together with proteins of interest. By examining overlap in expression, this workflow allows for exploring protein localization in structures that are challenging to distinguish based on histological examination only. This facilitates automated image analysis efforts, and minimizes the subjectivity introduced during manual annotation7,8.

Interestingly, when comparing the FE scores with single cell type specificity in the HPA, a dataset based on single-cell transcriptomics9, as many as 441 proteins with FE2-5 showed an elevated expression in ciliated cells. This suggests that further mapping of these cells is an important contribution towards understanding protein function. Here, we therefore seek to further explore the cilium (plural cilia), a filamentous hair-like cellular membrane protrusion, whose assembly (ciliogenesis) and structure requires at least 2,000 genes10. Cilia are canonically distinguished into two types: primary cilia and motile cilia. Motile cilia are present on epithelia of different organs where they aid in transporting cells, fluids, particles or mucus, and are crucial for e.g. reproduction, respiration, L/R body asymmetry, or brain development and function11. Defects in genes related to motile cilia have been linked to motile ciliopathies, contributing to a large spectrum of diseases in various tissues, including chronic respiratory problems, infertility, and hydrocephalus12. The most common ciliopathy, primary ciliary dyskinesia (PCD), is characterized by impaired ciliary movement. PCD is usually caused by autosomal recessive mutations, but also by autosomal dominant and even X-linked mutations in rare cases13,14.

Despite advancements in understanding PCD in the last 20 years, its diagnosis remains challenging, and effective treatments are lacking15. Studying motile cilia genes can provide us with fundamental insights into ciliary function and structure, being a first step towards understanding the underlying molecular mechanisms of ciliopathies (e.g., impaired mucociliary clearance in the airways)16. Structurally, the cilium comprises several distinct regions: the basal body, containing a modified centriole anchoring the cilium to the cell; the ciliary rootlet, a fibrous structure anchoring the basal body to the cell and it’s cytoskeleton for additional support; the transition zone, a specialized gateway at the ciliary base (between basal body and cilium) that regulates protein trafficking and acts as a diffusion barrier; and the axoneme, the core ciliary skeleton that extends from the basal body, traverses the transition zone and that is composed of microtubule doublets and associated motor proteins that mediate ciliary movement11,17. Given their small size, close spatial proximity, and dynamic nature, these compartments are best studied for their architecture using imaging and microscopy-based methods18. While electron microscopy and Cryo-ET have been instrumental in profiling ciliary ultrastructure, these techniques primarily reveal stable structural components like the axoneme, basal body, and transition zone, but often fail to preserve and visualize soluble or transiently associated proteins. Non-structural proteins yet remain underrepresented in structural studies. To our knowledge, no previous mIHC approach exists that enables comprehensive characterization of both structural and non-structural proteins in the different sub-compartments of ciliated cells at a high resolution. Therefore, we developed a medium-plex mIHC panel specifically designed to map cells with motile cilia.

With this background, the aim of the study was to develop a novel high-throughput workflow for quantifying cilia proteins at subcellular resolution using mIHC and automated image analysis, a first step towards understanding protein function. Based on signal overlap with a panel of marker proteins outlining the cilium, the transition zone, the rootlet, the cytoplasm, and the nucleus of ciliated cells, the spatial subcellular localization of 187 motile cilia proteins was mapped across five different tissue types from both respiratory (nasopharynx and bronchus) and reproductive tracts (fallopian tube, endometrium and cervix). The detailed information at the tissue, cellular, and subcellular levels offered new insights into the presumed functions of these proteins in motile cilia. Moreover, our approach, combining spatial proteomics with automated image analysis enables detailed spatial mapping at the subcellular level, offering a valuable resource for future large-scale studies of cilia biology in health and disease. The resulting images can be explored in the open access Human Protein Atlas (www.proteinatlas.org).

## Materials and Methods

### Ethics and tissue microarray design

Human tissue samples from five tissue types were obtained from Uppsala University Hospital, Sweden, and collected within the Uppsala Biobank organization. All samples were anonymized in accordance with the approval and advisory report of the Uppsala Ethical Review Board (ref nos. 2002-577, 2005-388, 2007-159). The tissue microarray (TMA) was generated as previously described19 and comprised five distinct tissues: fallopian tube, endometrium, cervix, nasopharynx, and bronchus (Figure 1B). Each tissue was represented by six one mm diameter cores, with duplicate cores from three individuals. Patient information is provided in Supplementary Table 1. The TMA was sectioned into 4 µm sections using a waterfall microtome (Microm H355S, ThermoFisher Scientific, Fremont, CA) and mounted onto SuperFrost Plus slides (Thermo Fisher Scientific, Fremont, CA). The slides were dried at room temperature (RT) overnight, baked at 50°C for 12–24 hours, cooled, and stored at RT until further use.

**Figure 1:**
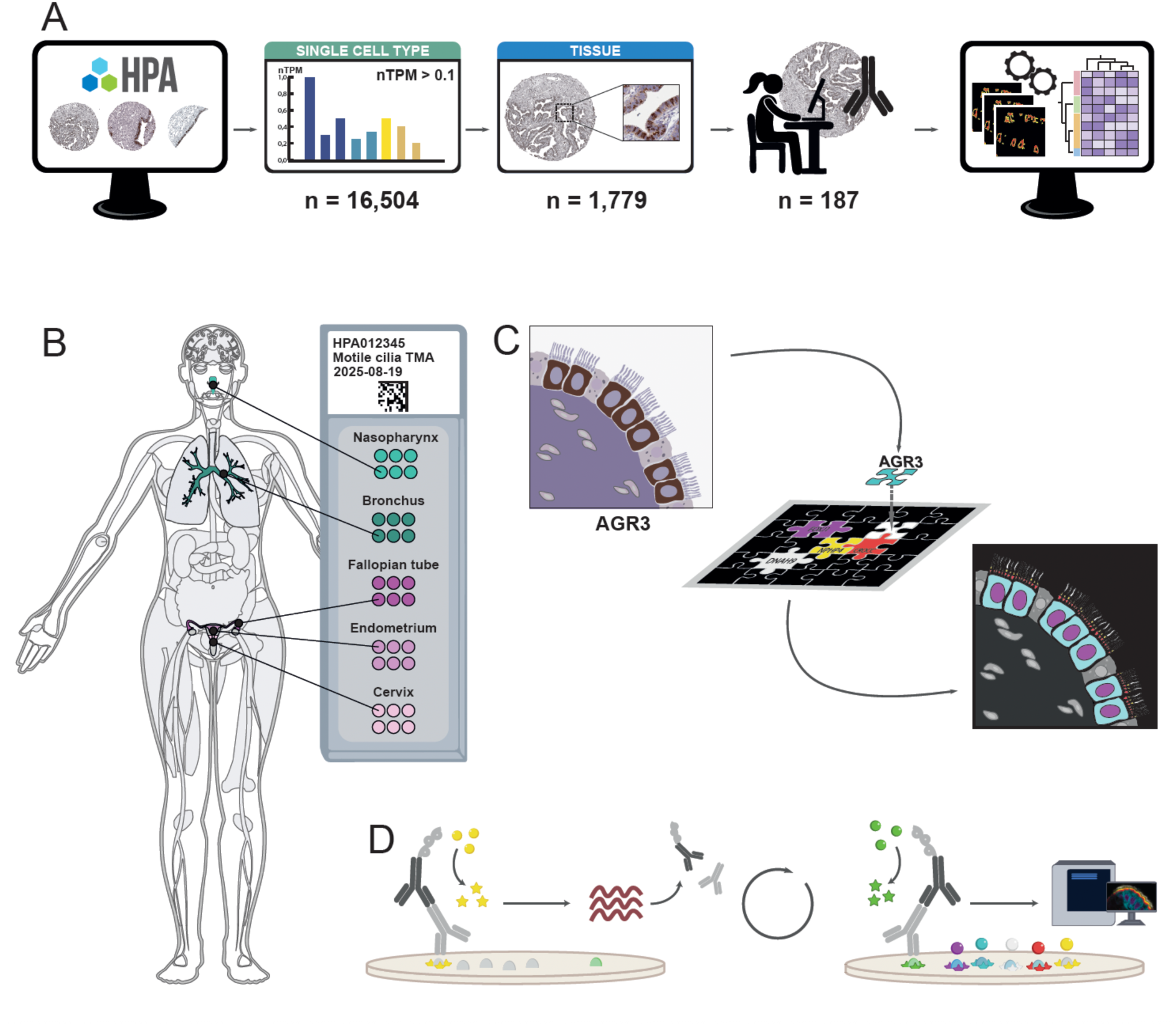
Experimental design. **A)** Selection of protein candidates from the Human Protein Atlas (www.proteinatlas.org) utilizing publicly available data from both the Single Cell Type and Tissue resources, allowing the identification of protein candidates expressed in ciliated cells with available reliably validated antibodies (n=187). The candidates were stained alongside the fixed antibody panel. Each acquired image was automatically analyzed to determine the expression profile, followed by clustering analysis. **B)** Five different tissues were collected and organized into a tissue microarray (TMA), consisting of two airway tissues (nasopharynx and bronchus) and three reproductive tissues (fallopian tube, endometrium, cervix). For each tissue, six cores from three patients were included. **C)** A multiplex immunohistochemistry (mIHC) antibody panel for ciliated cells was developed based on a literature review and existing IHC data in the HPA Tissue resource, here exemplified by the protein AGR3, showing staining in the cytoplasm of ciliated cells. The antibodies targeted different subcellular compartments of ciliated cells: nucleus (magenta), cytoplasm (cyan), rootlet (RL, red), transition zone (TZ, yellow), and cilium (CL, white). **D)** Schematic representation of the mIHC workflow, with the first and last cycles illustrated. The method relies on iterative cycles, each separated by heat-induced antibody stripping.

### Multiplex panel development

A fixed 5-plex panel was built to target five regions in ciliated cells: the cilium (CL), transition zone (TZ), rootlet (RL), cytoplasm, and nucleus. Protein markers and corresponding antibodies have been selected based on i) literature search (e.g. UniProtKB and scientific publications), ii) antibody IHC reliability score on HPA20 (see next section on Candidate selection), iii) expected IHC staining patterns in ciliated cells in nasopharynx, bronchus, fallopian tube (FT), endometrium, and cervix, and iv) compatibility with the multiplex immunohistochemistry (mIHC) panel and workflow. Each antibody was first tested with a single-plex run, i.e. one antibody at a time, to ensure that it worked equally well with the OPAL detection system, generating the same staining pattern as with regular IHC. Then, each antibody was tested in each position of the mIHC workflow to find the best location for each antibody to ensure a stable panel (Supplementary Table 2). Finally, when the best position for each antibody was selected, all five panel markers were stained simultaneously in a 5-plex staining, to confirm non-overlapping signals. To further validate the three markers targeting the cilium structure, confocal microscopy at 63X was conducted using Leica STELLARIS 5 (Leica Microsystems, Illinois, US).

### Candidate selection

Protein candidates and corresponding antibodies to be included in the present investigation were selected based on publicly available data in the HPA, including scRNA-seq data, IHC antibody reliability, and IHC staining pattern in human tissues. First, human scRNA-seq data from ciliated cells was used. This cell type in the HPA Single Cell Type Resource9 is based on external datasets from bronchus, lung, endometrium and fallopian tube. The data is presented as normalized transcripts per million (nTPM), showing the average expression of the cells in all cell type clusters from these four tissue types that represent ciliated cells. In this study, genes showing an expression level nTPM ≥ 0.1 were retained. Next, the IHC reliability score on HPA was considered, that evaluates the expected staining pattern in the entire human body taking into consideration positive and negative controls, literature, corresponding mRNA levels and similarity between multiple antibodies targeting the same protein. This stringent antibody validation pipeline is divided into four categories – Enhanced, Supported, Approved and Uncertain20. In this study, we restricted the selection to only antibodies with Enhanced, Supported or Approved reliability score. From this list, existing IHC annotation data from fallopian tube, bronchus and nasopharynx was considered, consisting of proteins that previously have been identified to stain specific subsets of epithelial cells in these tissues in the standard HPA pipeline. Finally, a manual curation was carried out based on IHC staining patterns in all tissues with motile cilia that were included in the present investigation, keeping only candidates with a distinct staining in ciliated cells, resulting in a refined list of candidate markers (Figure 1A).

### Slide pretreatment

To remove paraffin and rehydrate tissue cores, a ST5010 Autostainer XL (Leica Biosystems, Baden-Württemberg, Germany) was used with the following protocol: Xylene (5 min), Xylene (5 min), Xylene (1 min), absolute ethanol (3 min), absolute ethanol (3 min), 96% ethanol (3 min), 96% ethanol + 0.3% H₂O₂ (5 min), 80% ethanol (3 min), and dH₂O (30 sec). The slides were then stored in deionized water until antigen retrieval.

Antigen retrieval was performed to expose protein epitopes for antibody binding. Slides were subjected to heat-induced epitope retrieval (HIER) in pH 6 buffer (Agilent Technologies Inc., Santa Clara, CA, USA) at 125°C for 4 min using a Biocare Medical Decloaking Chamber™ PLUS (DC2008INTL). After cooling to 90°C, the slides were rinsed in deionized water (1–2 min) and transferred to TBS-Tween wash buffer (Thermo Fisher Scientific, TA-999-TT, Waltham, MA, USA) until further processing. To reduce autofluorescence, slides were incubated in a bleaching buffer containing 1.5% hydrogen peroxide, 0.2 M glycine, and 1X TBS-Tween (Thermo Fisher Scientific, TA-999-TT) in 50-mL Falcon tubes. Tubes were rotated under an overhead LED light at RT on a bench for 1 hour using a Stuart SRT9D roller mixer. Slides were subsequently washed and stored in TBS-Tween until further use.

### Multiplex immunofluorescence

For multiplex immunofluorescence (mIF) staining, one slide per candidate protein was used. Each of the six staining cycles proceeded as follows: Tissue sections were blocked with UV Blocking Buffer (10 min) (UltraVision LP HRP kit, Epredia, Kalamazoo, MI), incubated with primary antibody (30 min), rinsed in TBS-Tween, incubated with HRP-conjugated secondary antibody (pre-conjugated ready-to-use) (10 min) (UltraVision LP HRP kit, Epredia), and washed again in TBS-Tween. Slides were then incubated with the cycle-specific OPAL^TM^ fluorophore (Akoya Biosciences, Marlborough, MA) for 10 min, rinsed, and subjected to HIER (90°C, pH 6 buffer, (Agilent Technologies Inc.) 20 min). The order of the Opal dyes and panel markers from first to last cycle was: OPAL690 for CROCC, OPAL620 for DNAH9, OPAL520 for candidate protein (changed for each slide), OPAL570 for AGR3, OPAL480 for NPHP4, and OPAL-DIG-780 for FOXJ1. After each cycle, slides were rinsed and stored in wash buffer. The final step involved incubating slides with OPAL 780 fluorophore-conjugated anti-DIG antibody (1:125 in Epredia™ Lab Vision™ Antibody Diluent OP Quanto, #TA-125-ADQ) for 1 hour, followed by staining with DAPI (1:1000, Invitrogen, D1306, Thermo Fisher Scientific) for 5 min. Slides were rinsed, mounted with Invitrogen ProLong™ Glass Antifade Mountant, covered, and cured overnight. Finally, slides were digitized at 40X magnification using PhenoImager (Akoya Biosciences).

### Image analysis

A manual quality control was performed on all tissue cores, ensuring representativity of the staining pattern, presence of ciliated cells and normal tissue histology. Tissue cores that did not pass this quality control were not included in the data analysis. Automated image analysis was conducted in ImageJ Fiji21 to generate quantitative data. TIFF image stacks were processed per TMA core (Supplementary Figure 1A). Each stack contained separate images for panel marker proteins, DAPI, and autofluorescence. Images were converted to 8-bit, and DAPI/autofluorescence images were discarded (Supplementary Figure 1B). Segmentation was performed using 3-level Multi Otsu thresholding22, where only the highest threshold level (brightest pixels) was used (Supplementary Figure 1C). If segmentation failed for any marker, the stack was excluded from further analysis. For each panel marker image, segmentation masks were generated (Supplementary Figure 1D), defining regions of interest (ROIs) corresponding to each marker and candidate protein (Supplementary Figure 1E). ROIs were used to extract pixel values from candidate protein images by measuring the pixel value within each ROI, including mean pixel intensity, maximum intensity, minimum intensity, and total ROI area per core.

### Data analysis

Data analysis was performed using R 4.4.2. Median intensity values were calculated per ROI (CL, TZ, RL, cytoplasm, nucleus) for each protein and tissue, identifying predominant colocalization patterns. Cores excluded during manual annotation were omitted from median calculations. Data were normalized via Min-Max scaling per protein and slide across all tissues, with values rescaled between 0 and 1. Heatmaps were generated based on image analysis results using the pheatmap package (v1.0.12). Clusters in the heatmaps were generated using default Euclidean clustering incorporated in the package. Gene Ontology (GO) enrichment analysis was conducted using clusterProfiler (v4.14.6) and the human gene reference database org.Hs.eg.db (v3.20.0), retrieved on 2025-02-25. Only GO terms with p.value < 0.05 were retained. The top three terms per cluster were selected based on the highest number of associated proteins. The Protein Evidence (PE) and Function Evidence (FE) scores for the candidate proteins were downloaded from the HPP portal23 on 2025/02/13.

## Results

### A high-throughput workflow to study subcellular location of proteins in ciliated cells

Utilizing the extensive datasets in the publicly available HPA database (www.proteinatlas.org)5, we identified a total of 16,504 genes in the Single Cell Type resource that were expressed in the ciliated cell cluster above the defined expression threshold. Among these, 10,197 proteins were targeted by at least one antibody that met the IHC data reliability criteria, out of which 1,779 proteins were previously identified as having differential expression pattern in selected tissues harboring motile ciliated cells. A final list of 187 candidates for further exploration was retained following a manual examination of the IHC staining pattern in ciliated cells (Figure 1A; Supplementary Table 3). A dedicated tissue microarray (TMA) was created, including five tissue types known to harbor motile ciliated cells in both the respiratory and reproductive tracts: nasopharynx (N), bronchus (B), fallopian tube (FT), endometrium (E), and cervix (C) (Figure 1B; Supplementary Table 1). To precisely determine the localization of candidate proteins using an iterative mIHC workflow, a fixed panel of five antibodies outlining different subcellular structures of motile ciliated cells was developed (Figure 1C). The panel was combined with one antibody for a new candidate protein to be studied per slide (Figure 1D). This design allowed for a standardized high-throughput characterization of subcellular protein localization across multiple tissue samples.

The fixed panel was designed to map protein localization to five distinct subcellular regions of ciliated cells defined by the cilium (CL), the transition zone (TZ), the rootlet (RL), the cytoplasm, and the nucleus (Figure 2A). The final panel was based on the following proteins: DNAH9 (Dynein Axonemal Heavy Chain 9), a component of the motile cilia axoneme, here used as a marker for CL24; NPHP4 (Nephrocystin 4) localized in the TZ^25–27^; CROCC (Rootletin), the main structural protein of the ciliary RL28; AGR3 (Anterior Gradient Protein 3), described in mice as essential for regulating ciliary beat frequency, here used as a marker for cytoplasm29; and the transcription factor FOXJ1 (Forkhead Box Protein J1), a key transcription factor for motile cilia formation, an established marker for ciliated cell nucleus11,30 (Figure 2B; Supplementary Table 2). To validate the antibodies targeting the ciliary regions (CL, TZ, and RL), we performed confocal microscopy at 63X magnification (Figure 2C). Given the complex structure of the cilium, no substantial overlap was observed between these three markers, reinforcing confidence in the panel. The final validated mIHC panel enabled the development of image analysis pipelines and the definition of regions of interests (ROIs) based on the signals obtained from each of the markers in the fixed panel, corresponding to the five distinct subcellular localizations in ciliated cells (Supplementary Figure 1E).

**Figure 2:**
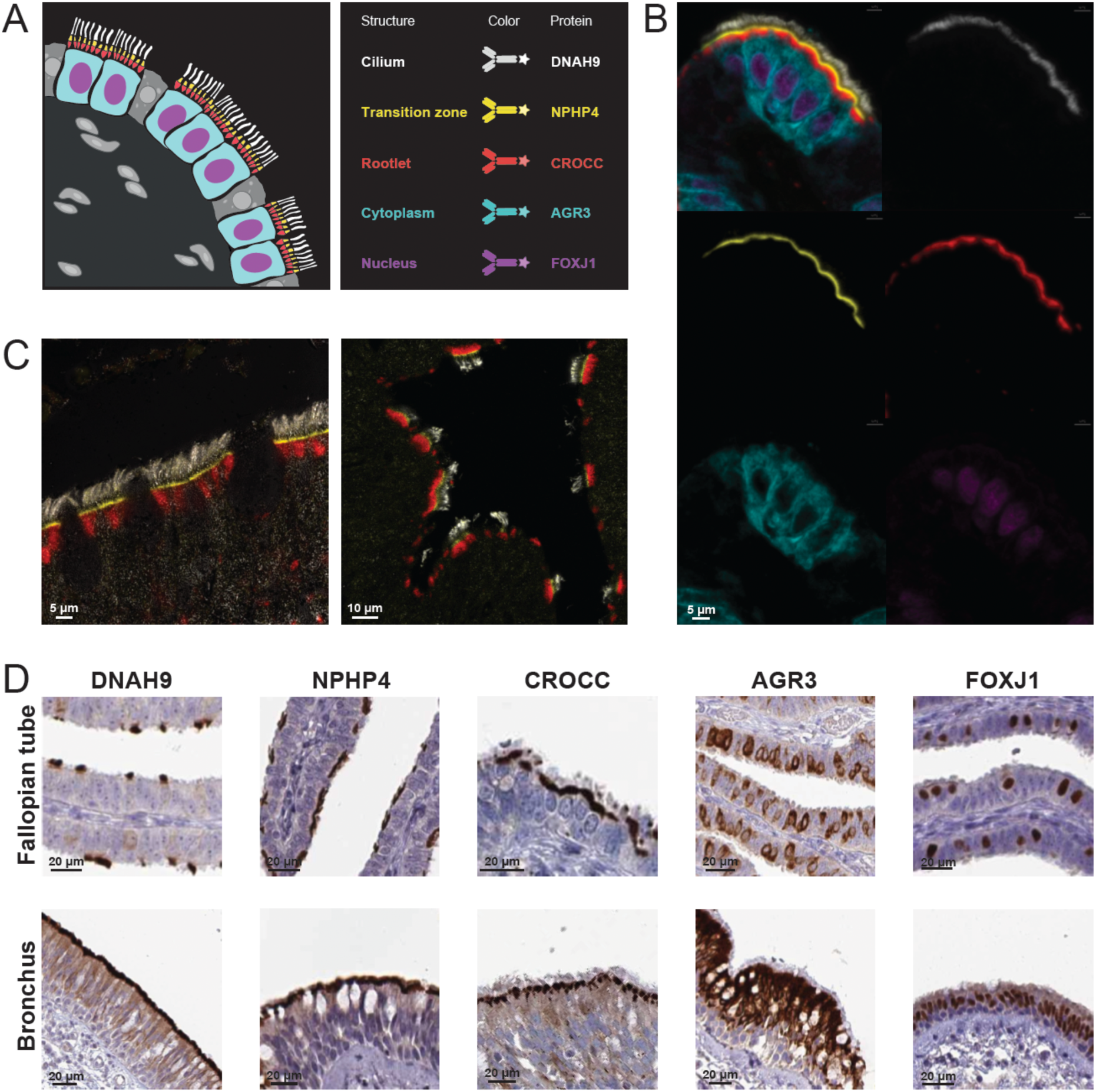
Validation and characterization of the fixed antibody panel. **A)** Schematic overview of the fixed panel used for mIHC stainings. The mIHC panel included five antibodies targeting proteins expressed in specific subcellular structures of ciliated cells: DNAH9 for the cilium (CL, white), NPHP4 for the transition zone (TZ, yellow), CROCC for the rootlet (RL, red), AGR3 for the cytoplasm (cyan), and FOXJ1 for the nucleus (magenta). **B)** Staining of the 5-plex fixed panel in ciliated cells of the fallopian tube. Top left: all five antibodies combined; top right: CL; middle left: TZ; middle right: RL; bottom left: cytoplasm; bottom right: nucleus. **C)** Confocal microscopy images. Confocal microscopy was used to validate the markers targeting ciliary structures. Non-overlapping signals representing the CL (white), TZ (yellow), and RL (red) are shown in the nasopharynx (left) and fallopian tube (right) at 63X resolution. **D)** IHC images from the HPA. Antibodies included in the panel were selected based on literature, as well as their staining patterns and reliability scores in the Tissue resource of the HPA.

### Hierarchical clustering and functional annotation of ciliated cells proteins

To address gaps in functional annotation, we integrated the FE scores defined by the HPP project20 with our 187 protein candidates (Supplementary Table 4) to assess the functional characterization of the proteins included in our study. Interestingly, just over a quarter (27%, n=51) of the candidate proteins stained alongside the panel fell into the FE1 category, indicating that at least one of their functions has been described. However, the majority (73%, n=136) remain poorly or partially characterized, with limited or no functional annotations (FE2 to FE5). To further examine the relationship between protein function and clustering, we analyzed the distribution of FE scores across clusters (Supplementary Figure 3). To provide a quantitative and systematic assessment of expression levels, we developed an automated image analysis workflow to quantify pixel intensity values for each analyzed protein and assess the subcellular localization of the staining patterns in relation to the fixed antibody panel (Supplementary Figure 1). To determine the overlap between each candidate protein and a specific area of the ciliated cells, the analysis method defined ROIs based on each panel marker individually. Pixel intensity values within these ROIs were then extracted from the images, allowing quantification of colocalization for each specific subcellular structure (Supplementary Figure 1). To account for multiple images per protein per ROI and per tissue, a median pixel intensity value was calculated per protein and tissue type for each ROI, generating an expression profile across the tissues. The resulting quantitative protein expression levels were subsequently used for hierarchical clustering, which identified seven distinct protein clusters (Figure 3B). Clustering was also performed on ROI per tissue type, revealing that CL and TZ regions consistently grouped together across all tissues. In contrast, RL, cytoplasm, and nucleus displayed more heterogeneous localization patterns, with the cytoplasm and RL exhibiting the highest variability across tissues. Gene ontology (GO) enrichment analysis was performed for each cluster across the three GO categories: biological process (BP), molecular function (MF) and cellular component (CC). This analysis aimed to validate the expression patterns across different subcellular localizations and provide insights into the potential functions of the protein within each cluster (Figure 3C; Supplementary Table 5).

**Figure 3:**
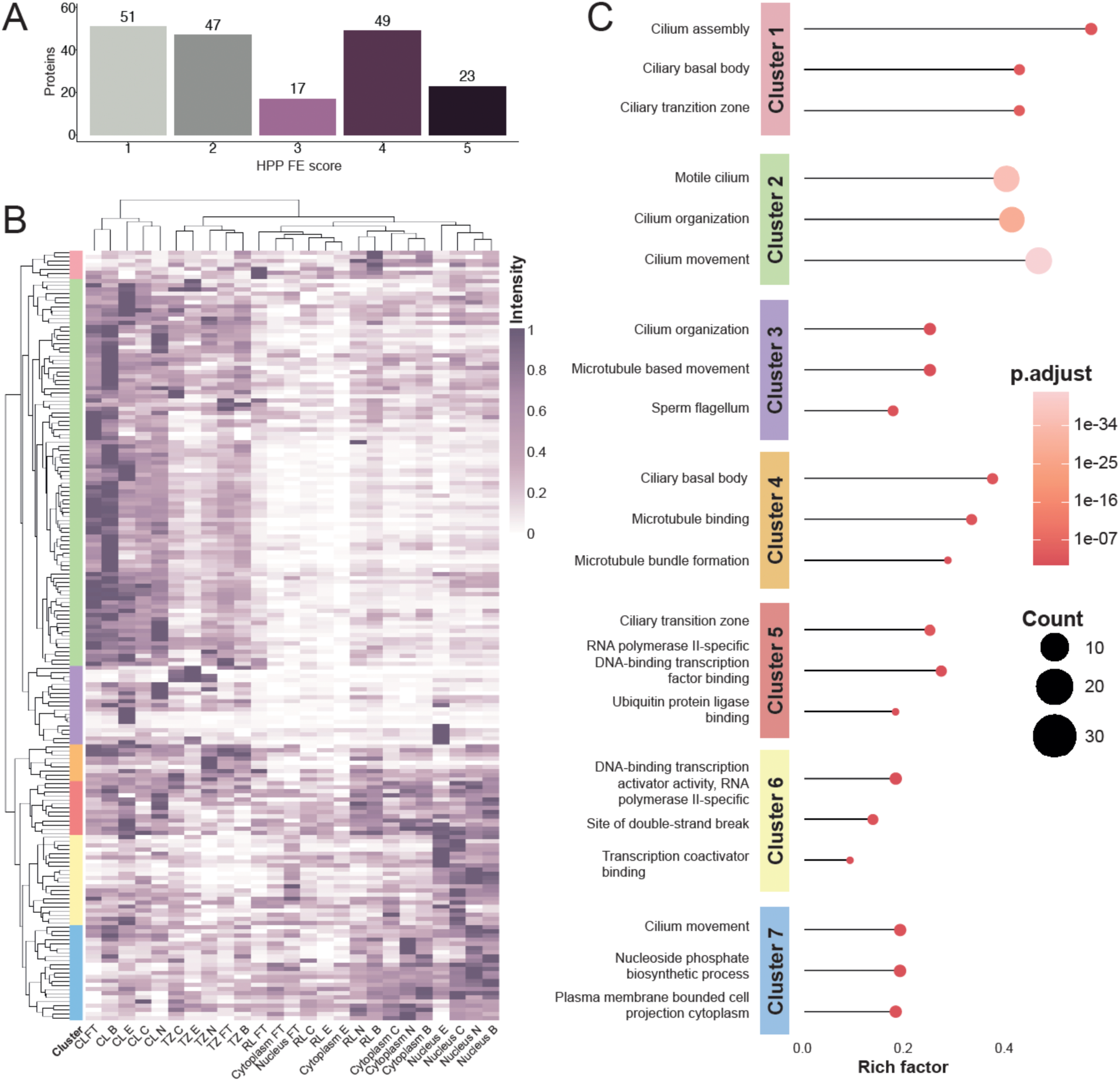
Functional evidence, Clustering Analysis, and Gene Ontology Enrichment. **A)** Bar plot showing the distribution of the studied protein between the different FE scores (FE1-5). **B)** A heatmap was generated based on expression levels obtained from image analysis for each candidate protein across different subcellular localizations and tissues (fallopian tube (FT), bronchus (B), endometrium (E), cervix (C), and nasopharynx (N). Hierarchical clustering was applied to both rows and columns. The columns were primarily clustered according to subcellular localization rather than tissue type, particularly for ciliary structures such as the cilium (CL) and transition zone (TZ). Row clustering identified seven distinct clusters based on the expression patterns of the 187 candidate proteins. **B)** Bubble plot showing the top three Gene Ontology (GO) terms of the three categories (BP, MF and CC) per cluster, with enrichment levels represented by the rich factor i.e. the proportion of input genes annotated to a specific term relative to the total genes associated with that term.

#### Cluster 1: Basal body and ciliary core

Cluster 1 (n = 7, primarily expressed in the RL, Figure 3B), was strongly linked to cilia, as indicated by its top GO terms: cilium assembly, ciliary basal body, and ciliary transition zone (Figure 3C). This suggests that proteins in this cluster play a central role in ciliary function. One notable example is MDM2, which localized to the RL of the cilia in all five tissues and showed a strong concordance between image observation and image analysis results (Figure 4A). Interestingly, none of the 65 GO terms associated with MDM2 in our analysis; including response to magnesium ion, cellular response to estrogen stimulus, positive regulation of intracellular protein transport, and negative regulation of cell cycle G1/S phase transition; are directly related to cilia or their structure. Although MDM2 is an FE1 protein with at least one well-characterized function, its role in ciliated cells remains unclear.

**Figure 4:**
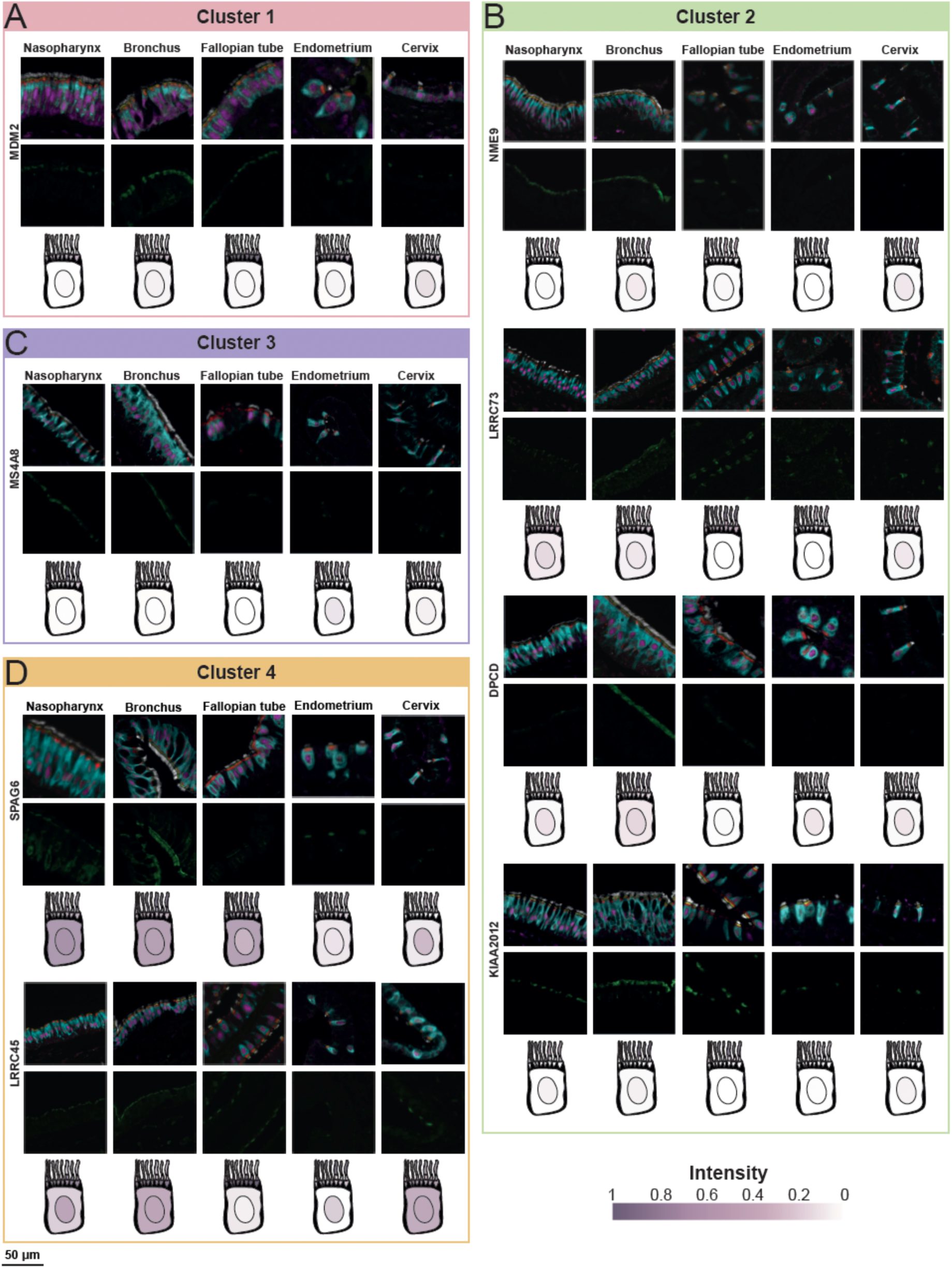
mIHC staining of representative proteins from each cluster across different tissues. The top row of images shows the composite staining, including the fixed panel and the candidate protein. The bottom row displays only the candidate protein (in green). Below the images, a schematic representation of a ciliated cell illustrates the expression levels obtained from image analysis for each subcellular localization. A) Cluster 1: MDM2 (RL) B) Cluster 2: NME9 (CL), LRRC73 (CL and RL), DPCD (CL), and KIAA2012 (CL). C) Cluster 3: MS4A8 (CL). D) Cluster 4: SPAG6 (whole cell in bronchus and nasopharynx; CL and TZ in endometrium and cervix) and LRRC45 (CL, TZ and RL).

#### Cluster 2: Axonemal and motile cilia proteins

Cluster 2 harbored the largest number of proteins (n = 94), with the top GO terms: motile cilium, cilium organization, and cilium movement, indicating a strong association with motile cilia specifically, rather than cilia in general (Figure 3C). The proteins in this cluster were primarily detected in CL across all five analyzed tissues (Figure 3B). One example is NME9 (NME/NM23 family member 9, also known as TXNDC6 or TXL-2) (Figure 4B). While NME9 is highly expressed in multiple tissues, with the highest mRNA level detected in the testis, previous studies have shown that it localizes to airway epithelial cilia as a microtubule binding protein31. Here, we demonstrated that NME9 is not restricted to the airway epithelium but is also present in the motile cilia of three female reproductive tissues, where it exhibited similar staining intensity. Another protein in Cluster 2 is LRRC73 (Leucine rich repeat containing 73) (Figure 4B). The mIHC analysis indicated that LRRC73 was equally expressed in CL and RL but absent in TZ, with a consistent pattern across all five tissues. However, no GO terms were identified for LRRC73 in our analysis, nor are functional annotations available in UniProt, and it is ranked as a FE5 protein (Supplementary Table 4 and 5). A third example is DPCD (Deleted in primary ciliary dyskinesia homolog) (Figure 4B), a protein implicated in the function or formation of ciliated cells. Its highest gene expression was observed in the testis, and previous studies have shown it is expressed in sperm32. Our analysis confirmed that DPCD was specifically detected in CL of motile cilia in all five tissues, with stronger expression in airway epithelial cilia compared to those in the female reproductive tract. In humans, DPCD interacts with RUVBL1 and RUVBL2, proteins known to be involved in dynein motor assembly, a crucial component for ciliary motility33,34. RUVBL1, which fell into Cluster 7, was detected in the cytoplasm and nucleus of ciliated cells but not in the ciliary structure (Figure 5C), in line with previous findings. GO terms identified for RUVBL1 include dynein axonemal particle, TFIID-class transcription factor complex binding, and ADP binding (Supplementary Table 4). Notably, *Ruvbl1* knockdown in mice leads to axoneme structural defects, further supporting its role as a ciliopathy-related protein essential for ciliary integrity35. Another protein found to be in Cluster 2 was ARMC2 (FE2), located in the cilium across all five tissues (Supplementary Figure 4). It has been shown that PCD caused by *ARMC2* mutations not only affects the reproductive system and abnormalities in sperm flagella36, but also results in an impaired airway system37. Here, spatial evidence is provided showing ARMC2 to be present in motile cilia in both female reproductive and airway epithelium (Supplementary Figure 4). Our analysis connected ARMC2 to multiple GO terms, such as cilium organization, cilium assembly, and spermatid development. Finally, KIAA2012, a poorly characterized protein with no known function (FE5) (Figure 4B; Supplementary Table 5) was also found in this cluster. Our data revealed that it localized to CL of motile cilia across all five tissues, with similar expression levels throughout, suggesting a conserved ciliary function.

**Figure 5:**
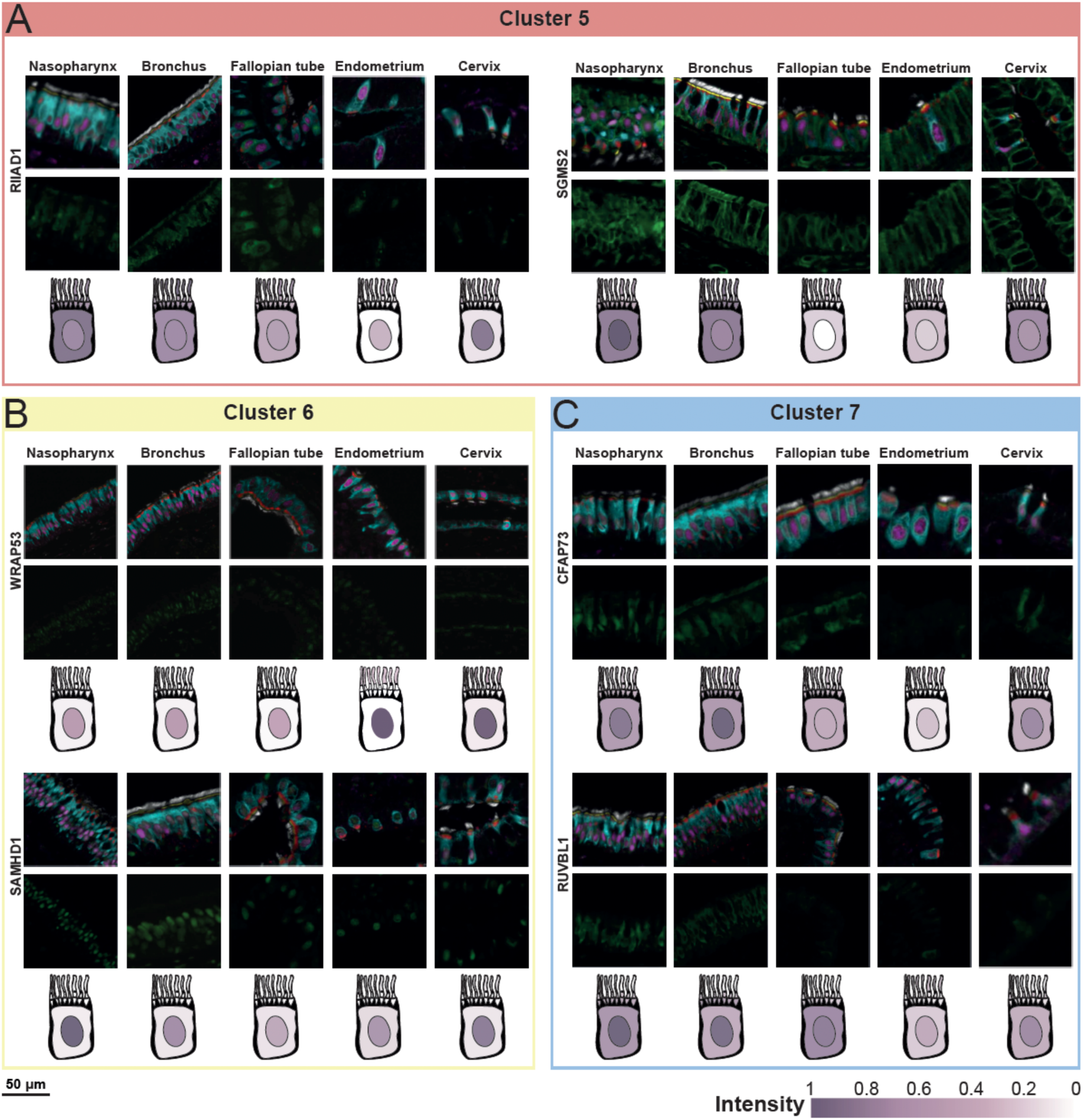
mIHC staining of representative proteins from each cluster across different tissues. The top row of images shows the composite staining, including the fixed panel and the candidate protein. The bottom row displays only the candidate protein (in green). Below the images, a schematic representation of a ciliated cell illustrates the expression levels obtained from image analysis, the darker the color, the higher the expression level. **A**) Cluster 5: RIIAD1 (CL, TZ, cytoplasm and nucleus) and SGMS2 (whole cell). **B**) Cluster 6: WRAP53 (nucleus) and SAMHD1 (nucleus). **C**) Cluster 7: CFAP73 (cytoplasm and nucleus) and RUBBL1 (cytoplasm and nucleus).

#### Cluster 3: Ciliary and flagellar motility proteins

The top GO terms for Cluster 3 (n= 19), were cilium organization, microtubule-based movement, and sperm flagellum (Figure 3C), highlighting its association with ciliary and flagellar structures. One notable protein in this cluster is MS4A8 (Membrane spanning 4-domains A8, also known as MS4A8b) (Figure 4C), a member of the MS4A protein family, that shares sequence and structural homology with the IgE receptor CD2038. Here, we demonstrated that MS4A8 is specifically localized to CL of motile ciliated cells across all five tissue types. MS4A8 has been ranked FE4 and interestingly also lacks previous evidence at the protein level (Supplementary Table 5). Our findings here provide spatial evidence of MS4A8 protein expression, offering a valuable insight into its potential role in motile cilia function.

#### Cluster 4: Dual localization in axoneme and transition zone

Cluster 4 (n = 9, with no FE1 proteins, Supplementary 3), presented a more complex expression pattern, with overrepresentation in both CL and TZ (Figure 3B). The top GO terms identified were ciliary basal body, microtubule binding, and microtubule bundle formation (Figure 3C), highlighting the potential structural roles of these proteins in cilia. One key protein in this cluster was SPAG6 (Sperm associated antigen 6) (Figure 4D), which is strongly linked to microtubule bundle formation, cilium organization, and cilium movement (Supplementary Table 5). SPAG6 is crucial for maintaining normal ciliary structure and function, and its loss has been associated with brain edema in mice39,40. While SPAG6 deficiency leads to male infertility, female mice remain fertile, although fertilization is delayed, suggesting an impact on the motile cilia in the oviduct41. In this study, SPAG6 showed widespread expression across the whole cell in bronchus and nasopharynx, while its localization was more restricted to CL and TZ in endometrium and cervix (Figure 4D). Another protein in this cluster is LRRC45 (Leucine rich repeat containing 45), which was primarily localized to CL, as well as TZ and RL (Figure 4D). Previously described in primary cilia, defects of LRRC45 are thought to alter ciliary length and beat frequency42,43. Here, we showed its presence and specific localization in motile cilia across both the respiratory and reproductive tract. However, our GO term analysis did not identify any functional annotation for LRRC45, and it is classified as FE5 (Supplementary Table 5), indicating that its precise role remains to be elucidated.

#### Cluster 5: Ciliated proteins with diverse localization

In Cluster 5 (n = 13), protein expression patterns were distributed across the entire ciliated cell (Figure 3B). The GO terms identified in this cluster were ciliary transition zone, RNA polymerase II-specific DNA-binding transcription factor binding, and Ubiquitin protein ligase binding, suggesting a broad functional involvement in cellular activity (Figure 3C). One example is RIIAD1 (Regulatory subunit of type II PKA R-subunit domain containing 1), which was predominantly localized in CL, TZ, cytoplasm and nucleus across all examined tissues (Figure 5A). Although RIIAD1 remains poorly characterized, studies have reported its upregulation in breast cancer tumors44,45. Notably, our GO term analysis did not identify any functional annotations for this protein, which is classified as a FE5 protein (Supplementary Table 4), suggesting its role in motile cilia should be examined. Another protein in this cluster is SGMS2 (Sphingomyelin synthase 2), which was detected in all targeted locations in the ciliated cells across all tissues, although significantly lower presence was observed in RL in the reproductive tissues (Figure 5A). SGMS2 is classified as a FE1 protein (Supplementary Table 4), indicating strong evidence for a molecular function. While our GO term analysis identified phosphotransferase activity for other substituted phosphate groups as a related term (Supplementary Table 5), additional annotations, including nucleoplasm, plasma membrane, and sphingomyelin synthase activity, suggest broader functional roles.

#### Cluster 6: Nuclear and cytoplasmic proteins

Cluster 6 (n = 22) consisted of proteins primarily localized to the cytoplasm and nucleus (Figure 3B), as supported by the top three GO terms: DNA-binding transcription activator activity, RNA polymerase II-specific, site of double-strand break, and transcription coactivator binding (Figure 3C). Notably, 45% of the proteins (n=10) in this cluster are classified as FE1, indicating strong evidence of the molecular function (Supplementary Figure 3). One key protein of this cluster is WRAP53 (WD repeat containing antisense to TP53) (Figure 5B), classified as FE1 (Supplementary Table 4). The GO term enrichment identified the site of double-strand break as a related function (Supplementary Table 5), aligning with its known role in DNA damage response. WRAP53 is recruited to the site of a double strand break (DSB), where it facilitates the recruitment of other DNA repair proteins46. Here, we showed that WRAP53 was predominantly expressed in the nucleus of ciliated cells across all tissues. Another protein in this cluster is SAMHD1 (SAM and HD domain containing deoxynucleoside triphosphate triphosphohydrolase 1) (Figure 5B), also classified as FE1 (Supplementary Table 4). Like WRAP53, SAMHD1 is associated with the GO term site of double-strand break (Supplementary Table 4). We revealed its nuclear localization in ciliated cells across all five tissues, reinforcing its potential role in DNA repair mechanisms.

#### Cluster 7: mixed localization with ciliary and cytoplasmic functions

The final cluster, Cluster 7 (n = 23), exhibited less specific localization patterns compared to other clusters, with proteins predominantly found in the cytoplasm and the nucleus (Figure 3B). GO term analysis identified key biological functions, including cilium movement, nucleoside phosphate biosynthetic process, and plasma membrane bounded cell projection cytoplasm (Figure 3C). Among the proteins in this cluster, RUVBL1 was previously mentioned with proteins from cluster 2 (Figure 5C) and classified as FE1 (Supplementary Table 4). Another protein in this cluster is CFAP73 (Cilia and flagella associated protein 73), classified as FE4 (Supplementary Table 4), which was primarily expressed in the cytoplasm and, to a lesser extent, in the nucleus across all tissues (Figure 5C). GO analysis linked CFAP73 to essential ciliary processes, including cilium movement, axonemal dynein complex assembly, and cilium organization (Supplementary Table 5), suggesting a role in ciliary structure and function.

## Discussion

This study introduces a comprehensive mIHC and automated image analysis workflow designed to investigate the subcellular localization of proteins in motile ciliated cells at a high resolution. To our knowledge, this is the first approach to combine high-resolution tissue imaging of motile ciliated cells with the scalability of high-throughput analysis of a large set of proteins across multiple tissue types using an in-house medium-plex panel tailored for this purpose. The unique panel and novel workflow were successfully used to spatially map 187 proteins in ciliated cells across five human tissues from both respiratory and female reproductive tracts. Protein localization was determined based on the overlap with a fixed reference panel outlining distinct subcellular compartments of motile ciliated cells. The validated panel enabled precise localization and quantification of each of the candidate proteins within five defined subcellular regions: the nucleus, cytoplasm, and three cilia-specific compartments: CL, TZ, and RL17.

The 187 candidate proteins studied in the present investigation were selected using two key resources of the Human Protein Atlas (HPA): i) the Single Cell Type resource, to validate the transcript-level expression in ciliated cells compared to other cell types in the studied tissues, and ii) the Tissue resource, to prioritize candidates with reliable antibodies showing positive IHC staining in ciliated cells in at least one of the five analyzed tissues included in the study. Using the cut-off threshold of 0.1 nTPM enabled the inclusion of a broad spectrum of proteins, encompassing both cilia specific markers and proteins with more ubiquitous expression profiles across different cell types (Supplementary Figure 2). As some proteins are broadly expressed across multiple cell types, immunostaining outside the ciliated cells population was occasionally observed and could be detected upon manual examination. The image analysis pipeline was however restricted to the predefined panel markers specifically targeting ciliated cells, and as a result quantitative data on protein expression in other cell types was not generated as part of this effort. One example of such a protein is GALNS (images not shown), that in addition to staining in ciliated cells also showed positivity in other epithelial cells and immune cells. Even if some proteins are not specific to ciliated cells from a body-wide perspective, we still believe that a detailed mapping of these proteins in ciliated cells aids in understanding their potential role in cilia function.

While most of the proteins encoded by the human genome have been credibly identified, many proteins still lack a specifically defined function based on experimental evidence. One of the major goals of the international consortium HPP is to gain a solid understanding of at least one function for every protein at the molecular level47. As a result, the HPP has introduced the definition of Function Evidence (FE) scores that build upon the functional information available in the UniProtKB, described as FE1-FE5 according to the level of evidence for a molecular function47. These new FE scores have been implemented for each protein encoded by the human genome. Interestingly, a majority (73%) of the candidate proteins characterized in our study showed FE2-FE5 scores, indicating that a full understanding of the role of these proteins in ciliated cells is still lacking. Our stringent validation pipeline and detailed data presenting a specific localization to ciliated cells at a subcellular resolution gives further insights into a presumed function of these proteins. The spatial localization data constitutes a starting point for further functional studies using e.g. model systems, to determine the roles of these proteins in each of the analyzed tissue types.

For further functional analysis, we also performed a hierarchical clustering analysis of the candidate proteins for each of the analyzed tissues using quantitative protein localization data generated by automated image analysis. The clustering analysis revealed that protein localization was largely conserved across tissues, particularly for CL and TZ, suggesting a shared spatial distribution for these proteins. Similarly, candidate proteins clustered based on their expression patterns, resulting in seven distinct expression groups. While some clusters exhibited clear localization patterns, such as cluster 2, with proteins mainly expressed in the cilium, or cluster 6, in the nucleus; others, like cluster 5, displayed a more dispersed expression profile. Nevertheless, the clustering analysis allowed us to group the proteins according to expression patterns, highlighting numerous poorly described proteins that shared a similar expression profile with well-described cilia markers, thus indicating a similar molecular function. We also performed a GO analysis for each of the expression clusters, which revealed a clear overrepresentation of cilia-associated functions, thus validating our strategy and providing an initial framework for a deeper understanding of protein function. It should however be noted that proteins localized to general cellular compartments such as the cytoplasm or nucleus may still exhibit cilia-specific roles. Given that GO annotations are derived from data across multiple cell types or model systems and not exclusively motile ciliated cells in the *in situ* context described here, definitive conclusions regarding specific functional or biological processes require further targeted investigation.

To confirm the validity of our approach, the list of candidates included several well-characterized proteins belonging to families with known functions in cilia, (such as cilia and flagella associated proteins (CFAPs), coiled-coil domain-containing proteins (CCDCs), dynein axonemal heavy chain proteins (DNAHs) and dynein axonemal intermediate chain proteins (DNAIs). We also shed light on several proteins with only limited evidence, such as seven different open reading frame proteins (C15orf48, C1orf87, C20orf8, C2orf50, C4orf47, C6orf132 and C7orf57) and other proteins with no previous description in the context of cilia biology, e.g. KIAA2012, RIIAD1 and MS4A8. Here we show for the first time that these proteins are expressed in specific subcellular compartments of motile ciliated cells. KIAA2012, a poorly described protein located in the cilium compartment (CL region) of motile ciliated cells in both respiratory and female reproductive tissues. Based on its subcellular location, it can be anticipated that this protein may contribute to cilia motility and thereby has the potential to cause primary ciliary dyskinesia (PCD) where the function of motile cilia is impaired, a syndrome causing a wide range of symptoms and diseases12. Another protein with limited characterization is RIIAD1, previously shown to be upregulated in breast cancer tumors, but without a known molecular function. The name of the protein suggests that it regulates a subunit of protein kinase A (PKA), involved in endocrine signaling and previously described to play a crucial role in regulating ciliary beating frequency48. Here, we show that this protein is generally expressed across the whole ciliated cell, both in the cell body and the cilium, in all five analyzed tissues. Interestingly, scRNA levels in other analyzed tissues in the Single Cell Type resource of the HPA show that expression is dominated in ciliated cells, followed by early spermatids, and cone photoreceptor cells. As mature sperm contain flagella, with a similar molecular function related to motility as ciliated cells49, and photoreceptor cells contain specialized sensory cilia50, the mRNA expression levels together with the spatial localization pattern suggests that this protein may be crucial for regulating and supporting key ciliary functions. Our study included numerous other proteins that lack a description of molecular function or were not previously described in the context of cilia biology, including MS4A8 which previously only has been described at the transcript level. Here, we could provide the first spatial evidence that this protein is expressed in the cilium of motile ciliated cells.

The selection of proteins included in the present investigation was mostly focusing on conserved motile ciliary proteins present in all the analyzed tissues, but we did identify several proteins whose expression pattern differed between respiratory and reproductive tissues. This shows that our workflow has the potential of cross-tissue comparison and that motile cilia may have different proteomes in different human tissue compartments, a finding that may explain phenotypic diversity in motile ciliopathy patients. Further studies using our novel strategy for mapping the entire motile cilia proteome in health and disease are expected to give unprecedented insights into the field of cilia biology.

Importantly, our method is facing limitations in resolution: Proteins detected in the TZ region might not necessarily be part of the TZ but localize close to it, at the rootlet or at the proximal cilium. Similarly, proteins that we discover in the CL region might not localize in the axoneme but in the cilioplasm or ciliary membrane. And we may detect proteins in the RL region that belong either to the RL or to broader structures close to the RL or in the basal body (BB). Nevertheless, our workflow provides a resolution not attainable by regular IHC, and a starting point for identifying subcellular expression patterns of ciliary proteins that can be validated by other molecular approaches.

In the rapidly expanding field of spatial biology, a multitude of methods focus on multiplexed imaging of proteins. Here, we developed a novel “medium-plex” workflow, centered around a fixed antibody panel outlining five different subcellular compartments of motile ciliated cells. While higher-plex methods enable the simultaneous analysis of multiple cell types and structures within a given tissue, our approach offers the advantage of subcellular-level resolution combined with high-throughput capacity, making it well suited for both proteome-wide investigation and clinical routine. This strategy provides a powerful foundation for comprehensive mapping of the motile cilia proteome in both health and disease. It represents an important step toward uncovering the molecular mechanisms driving ciliopathy and paves the way for future applications in precision diagnostics and personalized medicine.

## ASSOCIATED CONTENT

The following files are available free of charge (separate file (s)).

**Supplementary Figure 1:** Image analysis workflow

**Supplementary Figure 2:** mRNA expression levels across different cell types based on scRNA sequencing data for the 187 candidates included in the study

**Supplementary Figure 3.** Bar plot showing the distribution of FE scores across each expression cluster.

**Supplementary Figure 4.** mIHC staining of ARMC2 in the cilium in different tissues.

**Supplementary Table 1:** Patient information for all tissue samples included in the TMA

**Supplementary Table 2:** Information for the antibodies included in the fixed panel

**Other Supplementary Materials for this manuscript include the following:**

**Supplementary Table 3:** Information about the antibodies used to map the 187 protein candidates included in the study (separate .xlsx file)

**Supplementary Table 4:** Protein Evidence (PE) and Function Evidence (FE) scores from the HPP for the 187 candidate proteins included in the study (separate .xlsx file)

**Supplementary Table 5:** Protein expression levels based on automated image analysis, and results based on gene ontology enrichment for the 187 protein candidates included in the study (separate .xlsx file)

## Author Contributions

Conceptualization: CL and LM; Data curation: FB and LM; Formal Analysis: FB and LM; Funding acquisition: CL and MU; Investigation: FH, LM, JNH, FB; Methodology: CL, FB, LM and FH; Project administration: CL and LM; Resources: CL and LM; Image analysis: FB, FH, LM; Supervision: CL, LM, FH; Validation: FB, LM, and CL; Visualization: FB and LM; Writing – original draft: FB, LM and CL; Writing – review & editing: FB, LM, CL, FH, JNH and MU. All authors have read and approved the final version of the manuscript.

## Notes

Competing interests: Authors declare that they have no competing interests.

## Data availability

Out of the 187 proteins, 175 were made available in HPA version 25 (v25.proteinatlas.org). The remaining will be available in a repository up on publication.

## Code availability

The script for image analysis (Macros) and data analysis (R) used for data analysis and visualization is available on GitHub: github.com/LindskogLab/Subcellular-map-of-proteins-in-ciliated-cells

## ACKNOWLEDGMENT

We thank the Biological Visualization platform BioVis at Uppsala university for their vast help in imaging and image analysis. We also thank the patients who donated tissues for HPA. Lastly, we thank the members of HPA supporting this project in any way, especially Jonas Gustavsson, Rutger Schutten, and Borbala Katona.

## ABBREVIATIONS

AI: artificial intelligence
B: bronchus
BB: basal body
BP: biological process
C: cervix
CC: cellular component
CL: cilium
DSB: double-strand break
E: endometrium
FE: function evidence
FT: fallopian tube
GO: gene ontology
HIER: heat-induced epitope retrieval
HPA: Human Protein Atlas
HPP: Human Proteome Project
HRP: horseradish peroxidase
IgE: immunoglobulin E
IHC: immunohistochemistry
MF: molecular function
mIHC: multiplex immunohistochemistry
N: nasopharynx
PCD: primary ciliary dyskinesia
PE: protein evidence
PKA: protein kinase A
QC: quality control
RL: rootlet
ROI: regions of interest
RT: room temperature
scRNA: single-cell RNA
TMA: tissue microarray
TZ: transition zone

## For TOC Only

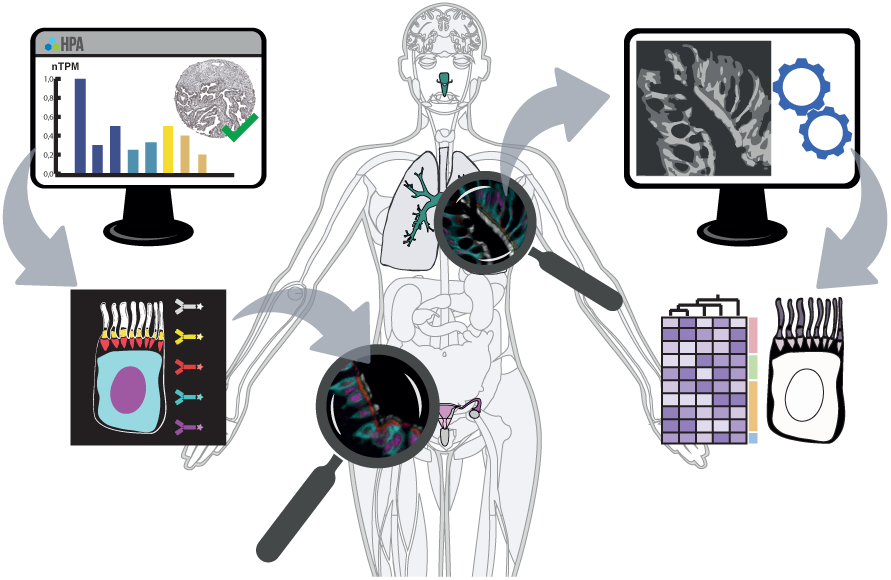

## REFERENCES

1. Hung, M.-C. & Link, W. Protein localization in disease and therapy. Journal of Cell Science 124, 3381–3392 (2011).

2. Thul, P. J. et al. A subcellular map of the human proteome. Science 356, eaal3321 (2017).

3. Adhikari, S. et al. A high-stringency blueprint of the human proteome. Nat Commun 11, 5301 (2020).

4. Omenn, G. S. et al. The 2024 Report on the Human Proteome from the HUPO Human Proteome Project. J Proteome Res 23, 5296–5311 (2024).

5. Uhlén, M. et al. Tissue-based map of the human proteome. Science 347, 1260419 (2015).

6. Method of the Year 2024: spatial proteomics. Nat Methods 21, 2195–2196 (2024).

7. Ghoshal, B., Hikmet, F., Pineau, C., Tucker, A. & Lindskog, C. DeepHistoClass: A Novel Strategy for Confident Classification of Immunohistochemistry Images Using Deep Learning. Mol Cell Proteomics 20, 100140 (2021).

8. Shariff, A., Kangas, J., Coelho, L. P., Quinn, S. & Murphy, R. F. Automated Image Analysis for High-Content Screening and Analysis. J Biomol Screen 15, 726–734 (2010).

9. Karlsson, M. et al. A single–cell type transcriptomics map of human tissues. Sci Adv 7, eabh2169 (2021).

10. Reiter, J. F. & Leroux, M. R. Genes and molecular pathways underpinning ciliopathies. Nat Rev Mol Cell Biol 18, 533–547 (2017).

11. Pedersen, L. B., Jurisch-Yaksi, N., Schmid, F. & Christensen, S. T. Cilia and Flagella. in Encyclopedia of Cell Biology (Second Edition) (eds Bradshaw, R. A., Hart, G. W. & Stahl, P. D.) 164–188 (Academic Press, Oxford, 2023). doi:10.1016/B978-0-12-821618-7.00209-1.

12. Wallmeier, J. et al. Motile ciliopathies. Nat Rev Dis Primers 6, 1–29 (2020).

13. Narayan, D. et al. Unusual inheritance of primary ciliary dyskinesia (Kartagener’s syndrome). J Med Genet 31, 493–496 (1994).

14. Krawczyñski, M. R., Dmeńska, H. & Witt, M. Apparent X-linked primary ciliary dyskinesia associated with retinitis pigmentosa and a hearing loss. J Appl Genet 45, 107–110 (2004).

15. Despotes, K. A., Zariwala, M. A., Davis, S. D. & Ferkol, T. W. Primary Ciliary Dyskinesia: A Clinical Review. Cells 13, 974 (2024).

16. Fliegauf, M., Benzing, T. & Omran, H. When cilia go bad: cilia defects and ciliopathies. Nat Rev Mol Cell Biol 8, 880–893 (2007).

17. Ishikawa, T. Structure of Motile Cilia. in Macromolecular Protein Complexes IV: Structure and Function (eds Harris, J. R. & Marles-Wright, J.) 471–494 (Springer International Publishing, Cham, 2022). doi:10.1007/978-3-031-00793-4_15.

18. Gopalakrishnan, J. et al. Emerging principles of primary cilia dynamics in controlling tissue organization and function. EMBO J 42, e113891 (2023).

19. Kampf, C., Olsson, I., Ryberg, U., Sjöstedt, E. & Pontén, F. Production of Tissue Microarrays, Immunohistochemistry Staining and Digitalization Within the Human Protein Atlas. J Vis Exp 3620 (2012) doi:10.3791/3620.

20. Sivertsson, Å. et al. Enhanced Validation of Antibodies Enables the Discovery of Missing Proteins. J Proteome Res 19, 4766–4781 (2020).

21. Schindelin, J., et al. Fiji: an open-source platform for biological-image analysis. Nat Methods 9, 676–682 (2012).

22. 廖 炳松, 陳 澤生 & 詹 寶珠. A Fast Algorithm for Multilevel Thresholding. Journal of Information Science and Engineering 17, 713–727 (2001).

23. HPP Portal. https://hppportal.net/.

24. Fliegauf, M. et al. Mislocalization of DNAH5 and DNAH9 in Respiratory Cells from Patients with Primary Ciliary Dyskinesia. Am J Respir Crit Care Med 171, 1343–1349 (2005).

25. Awata, J. et al. NPHP4 controls ciliary trafficking of membrane proteins and large soluble proteins at the transition zone. J Cell Sci 127, 4714–4727 (2014).

26. Fliegauf, M. et al. Nephrocystin Specifically Localizes to the Transition Zone of Renal and Respiratory Cilia and Photoreceptor Connecting Cilia. Journal of the American Society of Nephrology 17, 2424 (2006).

27. Czarnecki, P. G. & Shah, J. V. The ciliary transition zone: From Morphology and Molecules to Medicine. Trends Cell Biol 22, 201–210 (2012).

28. Nechipurenko, I. V. et al. A conserved role for Girdin in basal body positioning and ciliogenesis. Dev Cell 38, 493–506 (2016).

29. Bonser, L. R. et al. The Endoplasmic Reticulum Resident Protein AGR3. Required for Regulation of Ciliary Beat Frequency in the Airway. Am J Respir Cell Mol Biol 53, 536–543 (2015).

30. Wallmeier, J. et al. De Novo Mutations in FOXJ1 Result in a Motile Ciliopathy with Hydrocephalus and Randomization of Left/Right Body Asymmetry. Am J Hum Genet 105, 1030–1039 (2019).

31. Sadek, C. M. et al. Characterization of Human Thioredoxin-like 2. J Biol Chem 278, 13133–13142 (2003).

32. Zariwala, M. et al. Investigation of the Possible Role of a Novel Gene, DPCD, in Primary Ciliary Dyskinesia. Am J Respir Cell Mol Biol 30, 428–434 (2004).

33. Liu, G., Wang, L. & Pan, J. Chlamydomonas WDR92 in association with R2TP-like complex and multiple DNAAFs to regulate ciliary dynein preassembly. J Mol Cell Biol 11, 770–780 (2018).

34. Li, Y. et al. Cotranslational molecular condensation of cochaperones and assembly factors facilitates axonemal dynein biogenesis. Proc Natl Acad Sci U S A 121, e2402818121.

35. Dafinger, C. et al. Targeted deletion of the AAA-ATPase Ruvbl1 in mice disrupts ciliary integrity and causes renal disease and hydrocephalus. Exp Mol Med 50, 1–17 (2018).

36. Coutton, C. et al. Bi-allelic Mutations in ARMC2 Lead to Severe Astheno-Teratozoospermia Due to Sperm Flagellum Malformations in Humans and Mice. Am J Hum Genet 104, 331–340 (2019).

37. Wu, B. et al. Broadening the ARMC2 mutational phenotype: linking multiple morphological abnormalities of the Flagella to Pulmonary Manifestations in Primary Ciliary Dyskinesia. Reprod Biol Endocrinol 23, 48 (2025).

38. Liang, Y. & Tedder, T. F. Identification of a CD20-, FcɛRIβ-, and HTm4-Related Gene Family: Sixteen New MS4A Family Members Expressed in Human and Mouse. Genomics 72, 119–127 (2001).

39. Li, W. et al. Sperm Associated Antigen 6 (SPAG6) Regulates Fibroblast Cell Growth, Morphology, Migration and Ciliogenesis. Sci Rep 5, 16506 (2015).

40. Teves, M. E. et al. Sperm-Associated Antigen 6 (SPAG6) Deficiency and Defects in Ciliogenesis and Cilia Function: Polarity, Density, and Beat. PLoS One 9, e107271 (2014).

41. Sapiro, R. et al. Male Infertility, Impaired Sperm Motility, and Hydrocephalus in Mice Deficient in Sperm-Associated Antigen 6. Mol Cell Biol 22, 6298–6305 (2002).

42. Kanie, T. et al. A hierarchical pathway for assembly of the distal appendages that organize primary cilia. eLife 14, e85999 (2025).

43. Radhakrishnan, P. et al. Biallelic Variants in Impair Ciliogenesis and Cause a Severe Neurological Disorder. Clinical Genetics 107, 311–322 (2025).

44. Liu, X., Peng, Y. & Wang, J. Integrative analysis of DNA methylation and gene expression profiles identified potential breast cancer-specific diagnostic markers. Bioscience Reports 40, BSR20201053 (2020).

45. Lu, C. et al. Identification of hub genes in AR-induced tamoxifen resistance in breast cancer based on weighted gene co-expression network analysis. Breast Cancer Res Treat 197, 71–82 (2023).

46. Henriksson, S. et al. The scaffold protein WRAP53β orchestrates the ubiquitin response critical for DNA double-strand break repair. Genes Dev 28, 2726–2738 (2014).

47. Legrain, P. et al. The Human Proteome Project: Current State and Future Direction. Mol Cell Proteomics 10, M111.009993 (2011).

48. Kobayashi, A. et al. The Increase in the Frequency and Amplitude of the Beating of Isolated Mouse Tracheal Cilia Reactivated by ATP and cAMP with Elevation in pH. Int J Mol Sci 25, 8138 (2024).

49. Rosenbaum, J. L. & Witman, G. B. Intraflagellar transport. Nat Rev Mol Cell Biol 3, 813–825 (2002).

50. Hildebrandt, F., Benzing, T. & Katsanis, N. Ciliopathies. N Engl J Med 364, 1533–1543 (2011).

