## Supplementary figures + supplementary table 1 and 2 for "A high-resolution subcellular map of proteins in cells with motile cilia"

Filippa Bertilsson *et al.*

**This PDF file includes:**

**Supplementary Figure 1:** Image analysis workflow
**Supplementary Figure 2:** mRNA expression levels across different cell types based on scRNA sequencing data for the 187 candidates included in the study
**Supplementary Figure 3.** Bar plot showing the distribution of FE scores across each expression **cluster.**
**Supplementary Figure 4.** mIHC staining of ARMC2 in the cilium in different tissues.
**Supplementary Table 1:** Patient information for all tissue samples included in the TMA
**Supplementary Table 2:** Information for the antibodies included in the fixed panel

**Other Supplementary Materials for this manuscript include the following:**
**Supplementary Table 3:** Information about the antibodies used to map the 187 protein candidates included in the study (separate .xlsx file)
**Supplementary Table 4:** Protein Evidence (PE) and Function Evidence (FE) scores from the HPP for the 187 candidate proteins included in the study (separate .xlsx file)
**Supplementary Table 5:** Protein expression levels based on automated image analysis, and results based on gene ontology enrichment for the 187 protein candidates included in the study (separate .xlsx file)

**Fig. S1.**

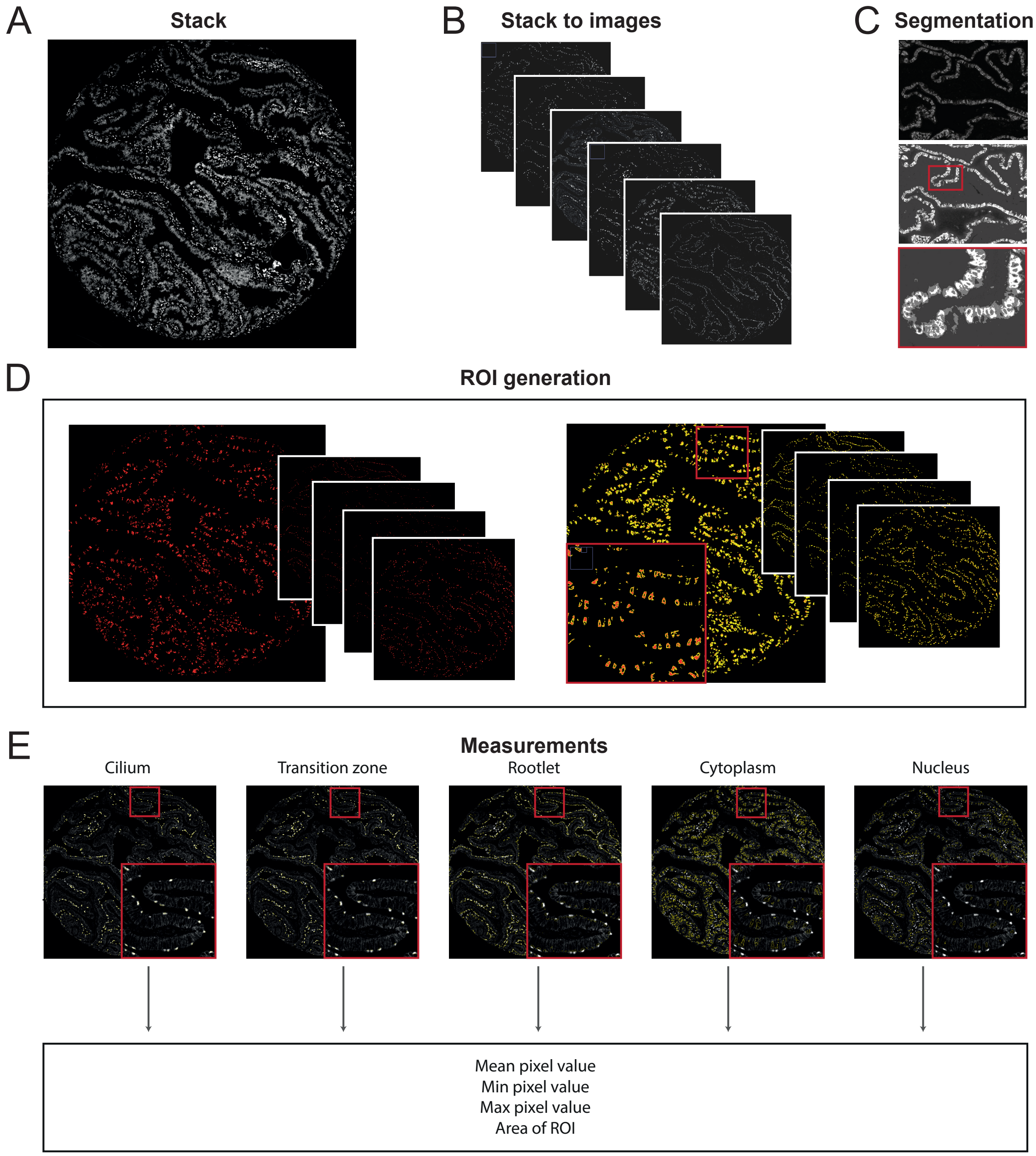


**Supplementary Figure 1: Image analysis workflow.** A) Input image. Images for the analysis were stacks, consisting of one image per protein plus one for DAPI and one for autofluorescence. B) Stack to images. Stack was separated to individual images and DAPI + autofluorescence image were discarded. C) Segmentation. Multi-Otsu segmentation was applied on each image with the panel markers, segmenting the pixels to three levels. The two lower levels with the lowest pixel values were discarded, and only the highest level (white) representing the positive pixels was used further. D) ROI generation. The segmentation (left, red) was used to generate regions of interests (ROIs) for each panel marker protein (left, yellow). E) Measurements. The ROIs were used to measure overlap on the candidate protein image. One ROI for one marker was used at a time to measure: mean pixel value, Min pixel value, Max pixel value, and Area of ROI. This was performed per core, and new ROIs were generated for each stack for each tissue and candidate protein. In this example, a core of fallopian tube was used.

**Fig. S2.**

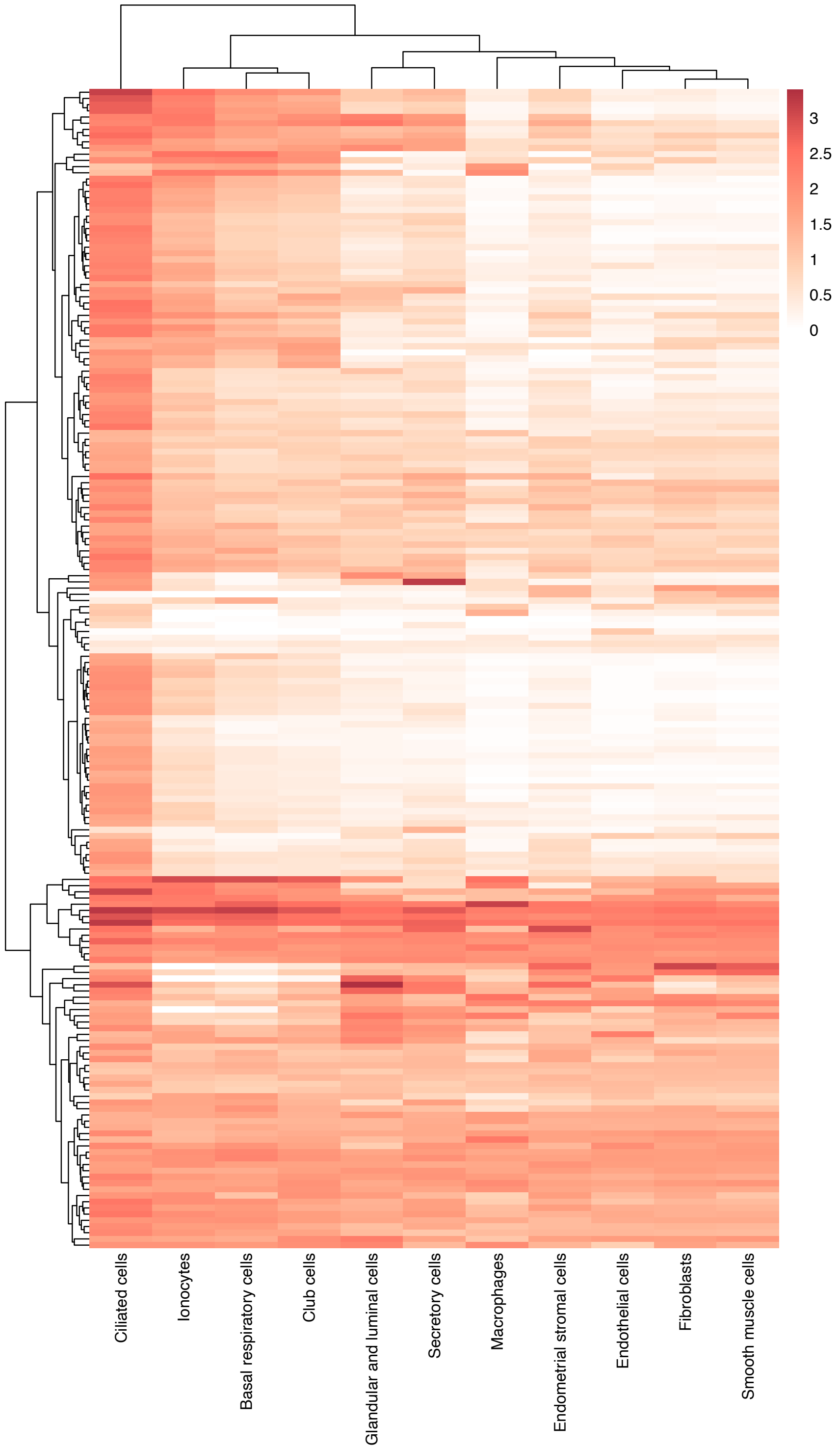


**Supplementary Figure 2: scRNA levels of studied proteins in ciliated cells compared to other cell types.** Normalized RNA levels (nTPM) on log10 scale in ciliated cells compared to other cell types found in the studied tissues for proteins incorporated in the study. Proteins with nTPM ≥ 0.1 was incorporated. Data from the Single cell resource on Human protein atlas (www.proteinatlas.org).

**Fig. S3.**

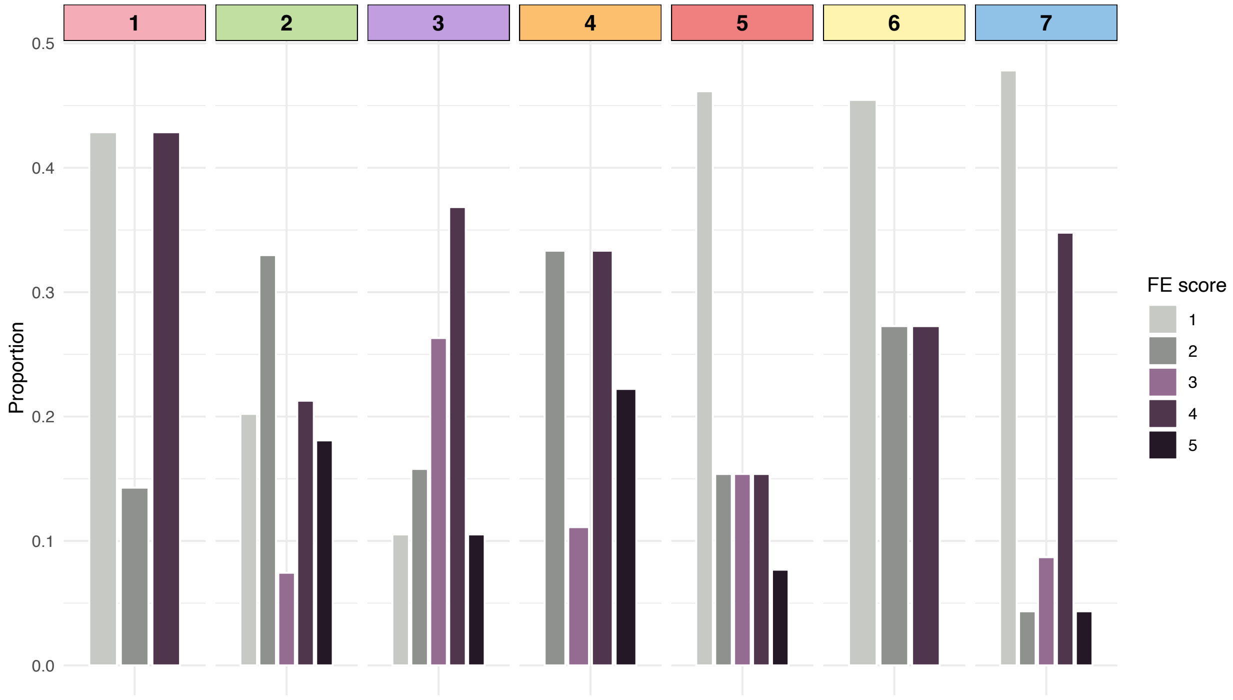


**Supplementary Figure 3. Functional evidence (FE) scores from The Human Proteome Project (HPP).** Distribution of the different FE scores (1-5) per protein cluster (1-7) identified based on protein expression. A FE1 score indicates that the protein has at least one well-described and experimentally validated function, while FE scores 2 to 5 reflect varying levels of uncertainty regarding the protein's molecular function.

Fig. S4.

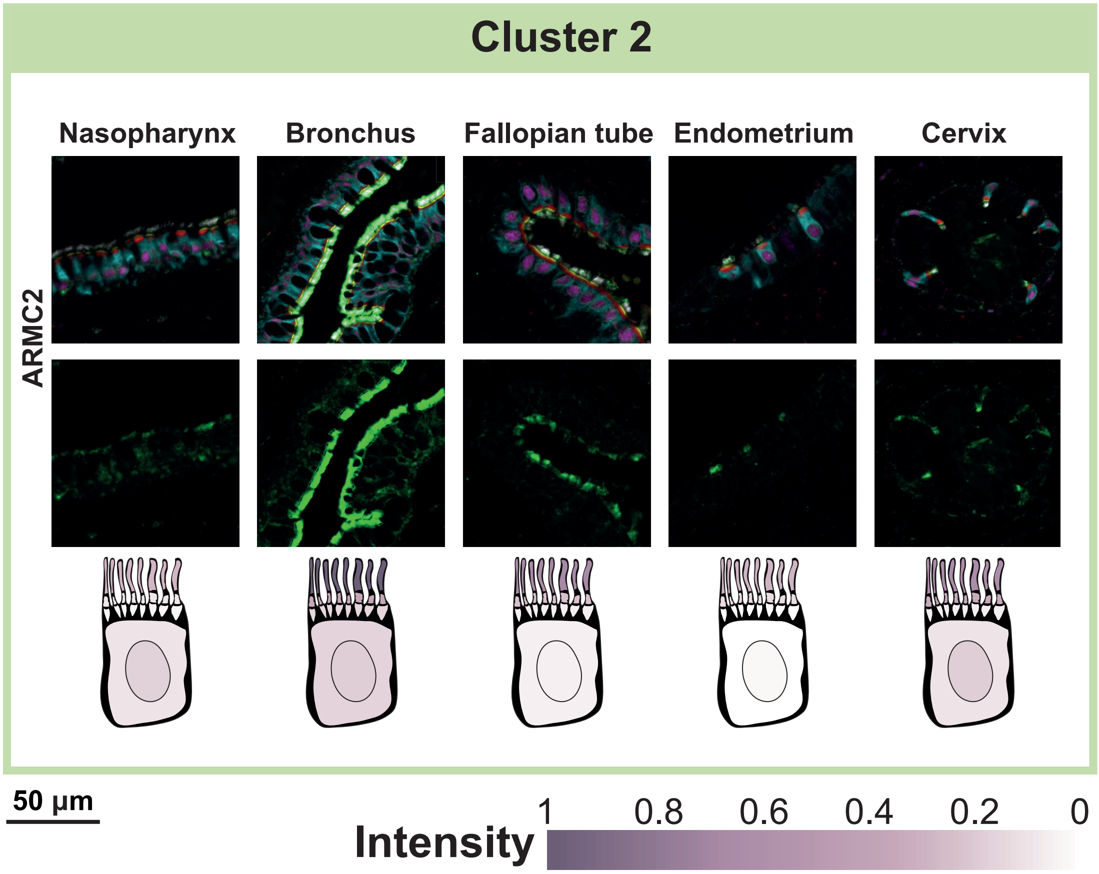


**Supplementary Figure 4. mIHC staining of ARMC2 in the cilium in different tissues**. The top row of images shows the composite staining, including the fixed panel and the candidate protein. The bottom row displays only the candidate protein (in green). Below the images, a schematic representation of a ciliated cell illustrates the expression levels obtained from image analysis for each subcellular localization.

**Table S1.**

| Tissue | Patient | Sex | Age | TMA |
|---|---|---|---|---|
| Fallopian tube | 2211.1 | Female | 25 | 1 |
| Fallopian tube | 2272.5 | Female | 37 | 1 |
| Fallopian tube | 1615.1 | Female | 47 | 1 |
| Cervix | 2079.1 | Female | 39 | 1 |
| Cervix | 777.2 | Female | 43 | 1 |
| Cervix | 52.2 | Female | 49 | 1 |
| Endometrium | 916.1 | Female | 41 | 1 |
| Endometrium | 2212.4 | Female | 43 | 1 |
| Endometrium | 915.3 | Female | 40 | 1 |
| Nasopharynx | 922.1 | Male | 13 | 1 |
| Nasopharynx | 1856.1 | Female | 81 | 1 |
| Nasopharynx | 1065.3 | Male | 48 | 1 |
| Bronchus | 2287.14 | Male | 18 | 1 |
| Bronchus | 2095.12 | Male | 58 | 1 |
| Bronchus | 2102.7 | Male | 82 | 1 |
| Fallopian tube | 2062.3 | Female | 43 | 2 |
| Fallopian tube | 1730.2 | Female | 49 | 2 |
| Fallopian tube | 2211.1 | Female | 25 | 2 |
| Cervix | 2079.1 | Female | 39 | 2 |
| Cervix | 9.4 | Female | 40 | 2 |
| Cervix | 52.2 | Female | 49 | 2 |
| Endometrium | 916.1 | Female | 41 | 2 |
| Endometrium | 2212.4 | Female | 43 | 2 |
| Endometrium | 375.1 | Female | 48 | 2 |
| Nasopharynx | 922.1 | Male | 13 | 2 |
| Nasopharynx | 1856.1 | Female | 81 | 2 |
| Nasopharynx | 1065.3 | Male | 48 | 2 |
| Bronchus | 2287.14 | Male | 18 | 2 |
| Bronchus | 2095.12 | Male | 58 | 2 |
| Bronchus | 2102.7 | Male | 82 | 2 |

**Supplementary Table 1: Patient information for all tissue samples included in the TMA.** Table of which patients were used for each tissue microarray (TMA) used in the study.

**Table S2.**

| Protein | Ensembl (v 109) | Antibody | Reliability score | Dilution | OPAL™ | Target structure |
|---|---|---|---|---|---|---|
| DNAH9 | ENSG00000007174 | HPA052641 | Enhanced | 600 | 620 | Cilium |
| NPHP4 | ENSG00000131697 | HPA065526 | Approved | 150 | 480 | Transition zone |
| CROCC | ENSG00000058453 | HPA021191 | Enhanced | 300 | 690 | Rootlet |
| AGR3 | ENSG00000173467 | HPA053942 | Enhanced | 150 | 570 | Cytoplasm |
| FOXJ1 | ENSG00000129654 | HPA005714 | Enhanced | 2000 | 780 | Nucleus |

**Supplementary Table 2: Information for the antibodies included in the fixed panel.** Information about the panel antibodies and their corresponding OPAL and target structure.
